# Glucose restriction reduces systemic, but not cardiac, inflammation during neonatal sepsis in preterm pigs

**DOI:** 10.64898/2026.10.14.759031

**Authors:** Ole Bæk, Björn Klabunde, Eman A. Mouhammad, Richard Dougthy, Páll S. Leifsson, Lisbeth H. Olsen, Jens P. Goetze, Thomas Thymann, Duc N. Nguyen

## Abstract

**Introduction:** Preterm infants are highly susceptible to sepsis and hyperglycemia may exacerbate inflammation and worsen outcomes. We examined how glucose provision influences systemic and cardiac responses during neonatal sepsis in preterm pigs.

**Methods:** Preterm piglets (90 % gestation) were infused with *Staphylococcus epidermidis* to induce sepsis. In Experiment 1, three ventilated and anesthetized piglets were continuously monitored for mean arterial pressure (MAP) and blood gases to characterize acute cardiovascular and metabolic deterioration. In Experiment 2, thirty-three piglets were randomized to standard (14 g/kg/day) or reduced (3.5 g/kg/day) glucose provision, with or without infection, and followed for 20h.

**Results:** Infection in Experiment 1 caused progressive hypotension, acidosis, and hypoxia, mirroring clinical neonatal sepsis. In Experiment 2, reduced glucose provision improved survival (77 % vs 46 %, p = 0.06) and attenuated systemic inflammation, with lower IL-6 and higher IL-10 concentrations. Infection induced mild cardiac neutrophil infiltration, without affecting plasma troponin T but clearly reduced cardiomyocyte mitotic density and triggered upregulation of inflammatory transcriptional pathways (NF-κB, JAK-STAT) and suppression of oxidative phosphorylation. In infected animals. glucose restriction preserved cardiomyocyte mitotic density but did not alter cardiac structure, injury markers or gene expression.

**Conclusion:** Neonatal infection induced circulatory failure, suppression of cardiomyocyte proliferation and upregulation of inflammatory genes in cardiac tissue, while reduced glucose provision mitigated systemic immune responses and preserved cardiomyocyte proliferation.

## Introduction

Around 15 million infants are born prematurely each year, and many face serious health challenges, including a high risk of infections[34]. These may progress to neonatal sepsis, which typically begins with an excessive inflammatory response that progress to organ dysfunction and circulatory collapse and remains a leading cause of death in this fragile population[39, 42]. Cardiac dysfunction is a frequent component of neonatal sepsis and strongly correlates with disease severity and poor outcomes[2, 39, 42, 52]

Whereas the increased risk of infections in preterm infants is traditionally attributed to immune immaturity and the need for prolonged medical support, such as mechanical ventilation and indwelling catheters[11, 21, 39, 43], the metabolic state of the preterm newborn can also contribute to poor infection outcomes[13, 14]. With limited fat, muscle, and glycogen reserves, newborns must allocate most energy to growth, leaving little for immune defense[14]. During infection, scarce energy may favour *disease tolerance*, maintaining health or tissue function despite infection, over energy-demanding *disease resistance* responses that aim to eliminate pathogens[4, 25, 38, 48]. However, when pathogen burdens exceed a threshold, resistance mechanisms are activated, requiring energy for immune cell proliferation, phagocytosis, and antimicrobial factors, generated primarily through aerobic glycolysis[24, 33]. This metabolic shift extends beyond immune cells, as we previously showed that high glucose administration during infection enhances hepatic glycolysis, drives systemic hyperinflammation, and reduces *disease tolerance*, increasing mortality and organ damage in preterm pigs with a serious bloodstream infection[6, 29, 53]. Conversely, enhanced hepatic oxidative phosphorylation supports systemic and local hepatic *disease tolerance* and improves survival[20, 50, 51].

Parenteral Nutrition (PN) is essential for preterm infants who are unable to tolerate full enteral feeding[44]. However, PN solutions typically contain high concentrations of glucose (around 10%), which frequently leads to hyperglycaemia[26]. Clinical studies have linked PN-associated hyperglycaemia to poorer outcomes during neonatal sepsis, suggesting that glucose provision may also affect organ function during infection[3, 41, 47]. However, it remains unknown how the highly metabolically active neonatal heart is affected by glucose availability during infectious episodes, and how cardiac energy metabolism itself is affected by neonatal infections. In newborns, cardiac energy is primarily generated through oxidative phosphorylation, with fatty acid β-oxidation providing approximately 70% of total energy[32]. During stress conditions like hypoxia, cardiac metabolism shifts towards glycolysis[8]. Such metabolic shifts, while initially adaptive, could become maladaptive during hyperglycaemia, potentially contributing to impaired mitochondrial function. In this study, we used a well-established preterm pig model of neonatal sepsis[1, 5] to address two main objectives. First, a characterization of cardiac dysfunction during infection. Second, investigating whether reducing glucose provision during infection alleviates cardiac metabolic and inflammatory disturbances. We hypothesized that neonatal infection would trigger marked cardiac inflammation and suppress mitochondrial metabolism and that reducing glucose provision would mitigate these effects.

## Methods

### Animal procedures

Experimental procedures and animal rearing have been explained in detail in previous publications[1]. The Danish Animal Experiments Inspectorate approved all experimental procedures (license number 2020-15-0201-00520), in accordance with EU Directive 2010/63, governing the use of animals in research. Briefly, piglets were delivered by caesarean section at 90% gestation (day 106/116) and after immediate resuscitation fitted with umbilical arterial catheters that allowed administration of PN and serial blood sampling. Then, pigs were placed in individual heated incubators with supplemental oxygen and experiments started ∼3 hours after birth. Experiments were initiated by infusion of live *Staphylococcus epidermidis* bacteria (SE, 5x10^9^ colony-forming units (CFU)/kg) and pigs were closely monitored for the next 20 hours or until humane endpoints for euthanasia were met. Humane endpoints were arterial blood pH of <7.1, severe lethargy, hypoperfusion, irregular respiration or spontaneous skin bleeding. Animals were deeply sedated (Zoletil, Denmark) prior to euthanasia by intracardiac injection of pentobarbital (140mg/kg, Euthasol vet, Denmark).

### Experiment 1: Intensive monitoring of cardiac function during neonatal sepsis

Three newborn preterm piglets were subjected to intravascular arterial blood pressure measurements and close monitoring of blood gas parameters. Catheters were inserted into the umbilical vein and artery immediately after birth. The piglets were placed under general anaesthesia, maintained via continuous infusion of a 1:1 solution of propofol (5 mg/mL, Propofol “B.Braun”, Germany) and isotonic saline through the umbilical vein catheter. Upon achieving full aesthetic depth, the piglets were placed in ventral recumbency, intubated with a 2.5–2.3 mm inner diameter cuffed tracheal tube, and connected to a ventilator (BabyLog VN500, Dräger, Germany) set to *Volume Guarantee* mode (PC-PSV) with the following baseline settings: Tidal Volume (VT) = 5 mL/kg; Frequency (f) = 30/min (inspiration time 0.6 s, expiration time 1.4 s); Inspired Oxygen Concentration (FiO2) = 21%; Maximum PIP = 25 cm H2O; and Maximum PEEP = 5 cm H2O. The parameter used to evaluate steady-state conditions was SpO2 ≥ 90%, and FiO2 was adjusted to support a high SpO2. To maintain the piglets’ body temperature, they were placed on a *Bair Hugger* (Solventun, USA) heating mattress during anaesthesia, with a rectal probe secured to the tail for continuous monitoring of body temperature. Systemic arterial blood pressure and heart rate were measured continuously via the umbilical artery. Blood gas measurements from arterial blood were likewise performed every 5-10 minutes (Radiometer, Denmark). The experiment was initiated by infusion of SE bacteria and pigs were closely monitored until humane endpoints were met. Pigs were kept on standard 10% glucose PN (14 g/kg/day) for the duration of the experiment.

### Experiment 2: Effects of differing glucose provision during neonatal sepsis

In *Experiment 2* a total of 33 preterm piglets were randomized to receive either standard glucose (10%, 14 g/kg/day, CON) or restricted glucose PN (2.5%, 3.5 g/kg/day, RES), followed by infusion with either control saline (CON) or live SE, resulting in four groups: CON-CON (n=3), RES-CON (n=4), CON-SE (n=13), and RES-SE (n=13). Animals were monitored until they reached humane endpoints or up to 20 hours post-infection. Blood samples were collected at 3, 6, and 12 hours post-infection and at euthanasia for blood gas analysis, haematology and measurement of blood bacterial burdens as well as plasma cytokine, troponin T and biochemical profiling After euthanasia, the heart was removed and weighed; the apex was snap-frozen for RNA sequencing, and transverse sections of both ventricles were fixed in formalin for histological analysis. *Experiment 2* was performed as a part of a larger experiment where main results have been reported elsewhere[20].

### Cardiac endpoints

Plasma levels of troponin T were measured using an automated COBAS platform (Roche Diagnostics, Germany). From the formalin-fixed transverse ventricular slice, full-thickness blocks of the left ventricular free wall (LV), interventricular septum (IVS) and right ventricular free wall (RV) were paraffin-embedded, sectioned at 3 µm, stained with H&E. Slides were microscopically inspected for signs of necrosis, cellular damage (hydropic degeneration) and neutrophil infiltration. To determine mitosis, images were analysed in QuPath v0.7.0 (Bankhead et al., 2017) after coding, in random order, by an observer blinded to group. In each region the myocardium was outlined as a region of interest (ROI), excluding a 500 µm edge band, folds, crush artefact, large vessels and epicardial fat. Nuclei within the ROI were detected on the haematoxylin channel with fixed, pilot-validated settings. Each ROI was screened at 20× on a 500 µm grid, and candidate figures were confirmed at 40×. A mitotic figure was defined as condensed chromosomes without a nuclear membrane, from prometaphase to telophase[9] while prophase, pyknotic and apoptotic nuclei were not counted. Figures were marked as point annotations and classed as cardiomyocyte when they lay within striated sarcoplasm aligned with the fibre axis, other cells were classified as noncardiomyocytes. Outcomes were cardiomyocyte mitotic density (figures/mm²) and mitotic index (figures/1,000 nuclei).

From snap frozen cardiac tissue, total RNA was isolated using the QIAGEN RNeasy Mini Kit (QIAGEN, Germany) following the manufacturer’s protocol. Briefly, 15–30 mg of tissue was minced on ice and homogenized in lysis buffer using the gentleMACS™ Dissociator (Miltenyi Biotec, Germany). Samples were centrifuged, and the supernatant was mixed with 70% ethanol before being loaded onto silica-based spin columns. After multiple wash steps and DNase treatment to remove genomic DNA, RNA was eluted in RNase-free water. RNA concentration and purity were assessed with a NanoDrop spectrophotometer (ND-1000, Thermo Scientific, USA), and samples were stored at –70°C before sequencing at a commercial partner. (Novogene Co, USA).

### Statistics

Statistics were performed in Stata 14.2 (StataCorp, USA). In *Experiment 1*, for correlations between clinical or blood gas measurements and time, or between mean arterial pressure and blood gas measurements, a repeated measures fixed-effects linear regression model with time as a continuous variable was used to account for within-subject variation. Model fit was assessed by comparing linear, quadratic, and non-linear specifications, and the model with the best fit was selected. The within-subject R² is reported as a measure of explained variance. In *Experiment 2*, to analyse changes in the outcome over time and evaluate effects of different glucose regimens or SE infection, we used a linear mixed-effects model with fixed effects for time, treatment, sex, birthweight, and litter. A random intercept for each animal was included to account for repeated measurements within individuals. To evaluate systemic disease tolerance and resistance states, reaction norm analyses were performed by simple linear regression where pH was plotted against blood bacterial density separately at each timepoint and within each treatment group, as described previously[50]. Slopes were compared between groups and P value of differences reported if significant.

For differential gene expression (DEG) analysis in cardiac tissue, statistical analysis, data handling and visualization was performed in R (version 4.5). DEGs included only protein-coding genes with ≥10 counts in at least four samples. DEGs were identified using *DESeq2* (v1.38.3) across two comparisons: RES-SE vs. CON-SE and SE vs. CON, adjusting for sex and litter. Fold changes were estimated with *lfcShrink*, and transcripts with FDR < 0.05 were considered significant. Volcano plots were generated with *EnhancedVolcano* (v1.14.0). Gene set enrichment analysis (GSEA) was performed using *clusterProfiler* (v4.0.2), ranking genes by log₂ fold change. Pathway annotations were based on Kyoto Encyclopedia of Genes and Genomes (KEGG) and Gene Ontology, Biological Process (*Sus scrofa*). Pathways with FDR < 0.05 were deemed significant; normalized enrichment scores indicated up- or down-regulation. Heatmaps of Z-score-scaled gene expression were produced using *ComplexHeatmap*.

## Results

### Characterization of cardiac responses to neonatal sepsis

In *Experiment 1*, the median survival time for infected piglets was 8.2 hours. Mean arterial pressure (MAP) remained stable at approximately 30 mmHg during the first four hours after infection but then declined steadily until humane endpoints for euthanasia were reached (Figure 1A). This drop in MAP was accompanied by a gradual compensatory increase in heart rate over the course of the infection (Figure 1B). The decline in blood pressure coincided with the development of metabolic acidosis and hypoxia, reflected by a decrease in blood oxygen saturation and pH along with a rise in lactate levels (Figure 1C–E). During the infection the blood glucose levels dropped steadily from around 7 to 5 mmol/l over the course of the experiment (Figure 1F). As such we noted a clear correlation between drop in arterial pressure and acceleration of the metabolic acidosis accompanying neonatal sepsis.

**Figure 1:**
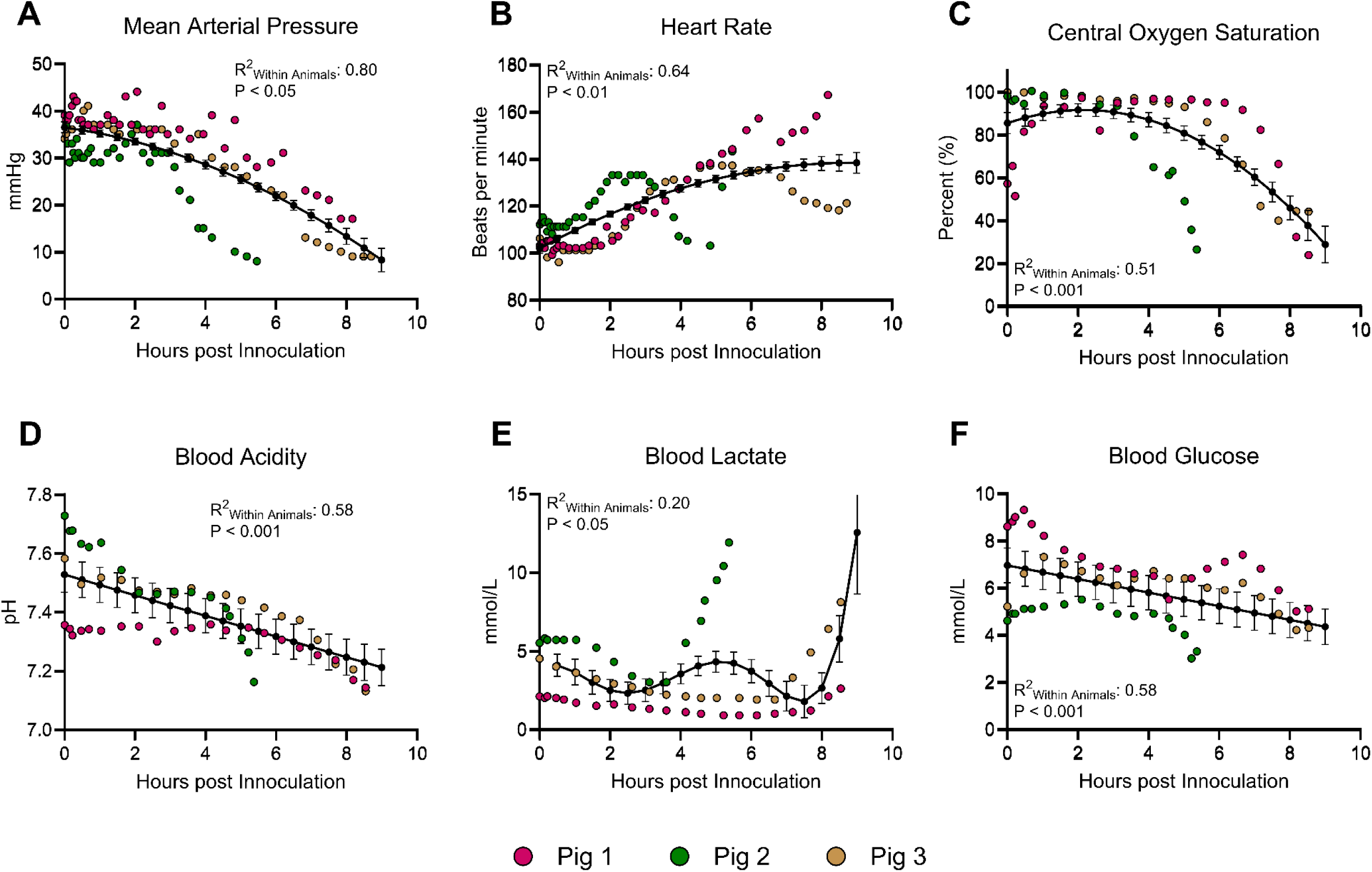
Cardiac and blood gas parameters in three anesthetized preterm pigs fitted with ventilators and infused with live *Staphylococcus epidermidis* (SE) bacteria. **A, B:** Mean arterial pressure (MAP) and heart rate measured by invasive monitoring. **C–F:** Blood gas parameters and blood metabolites. Data are shown as individual values with model-derived means and corresponding standard errors. *P* values indicate changes over time, and *R*² values represent model fit.

### Reduced glucose provision decreases clinical severity and inflammatory responses during infection while improving survival

In *Experiment 2*, infected piglets receiving 10% glucose, only 46% (6 of 13) survived until the end of the experiment, compared to 77% (10 of 13) in the RES-SE group. Although this difference did not reach statistical significance (Figure 2A, P = 0.06), it suggests a survival benefit with reduced glucose provision, which we have shown before[6, 29, 51]. However, there were no statistical differences in blood pH, oxygenation, or lactate levels between groups, although piglets in the RES-SE group showed a trend toward reduced metabolic acidosis (Figure 2B,C). This was observed despite clear differences in blood glucose concentrations between the two groups (Figure 2D). As expected, SE infection alone induced marked metabolic acidosis, characterized by low blood pH and elevated lactate levels (Figure 2B,C).

**Figure 2.**
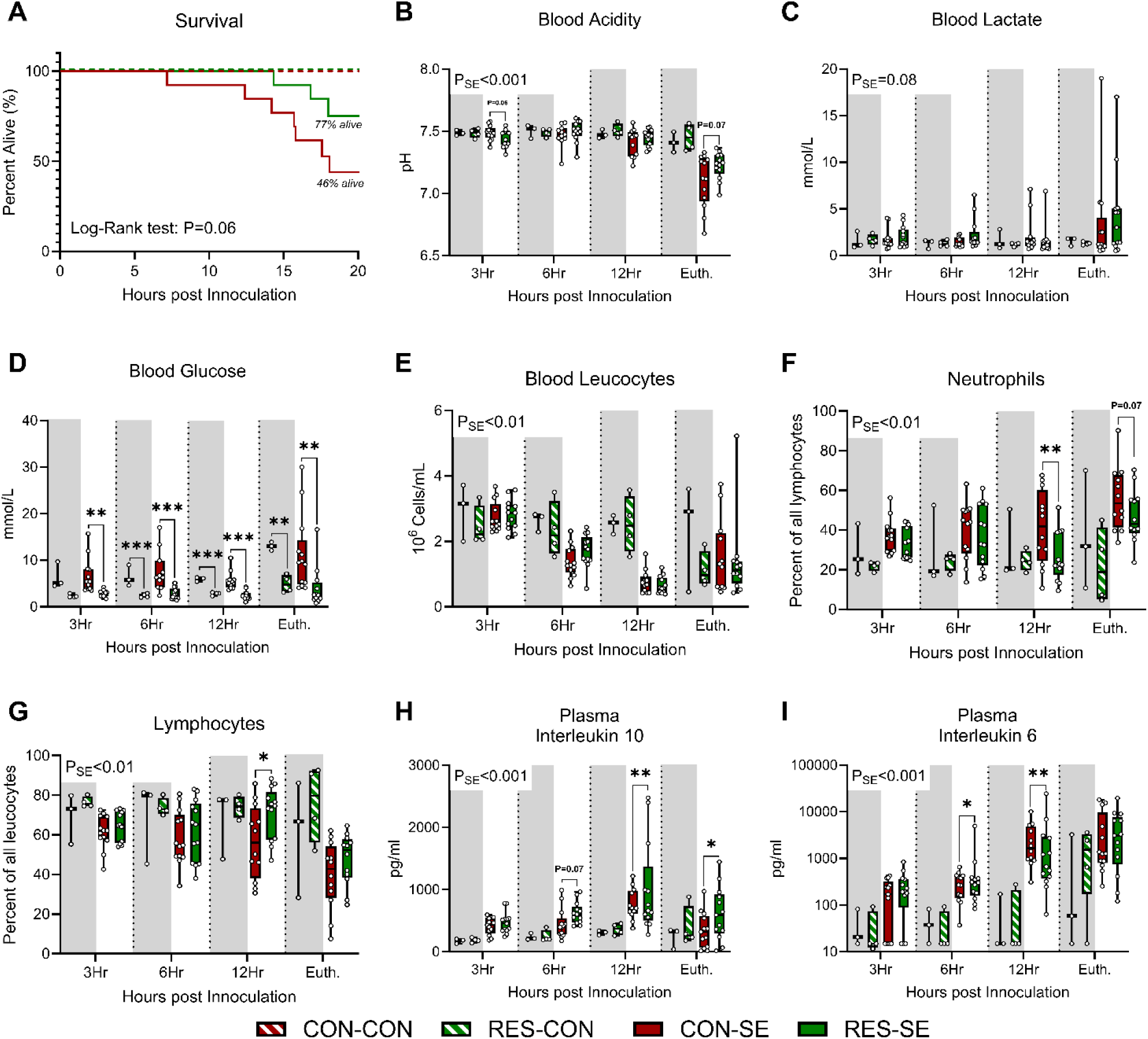
Clinical and paraclinical responses in preterm pigs nourished with either standard (CON) or restricted (RES) glucose parenteral nutrition (PN) and infused with either *Staphylococcus epidermidis* (SE) bacteria or saline (CON). **A:** Survival curves with corresponding log-rank test. **B–D:** Blood gas parameters. **E–G:** Hematological parameters. **H, I:** Plasma cytokine levels. Results from mixed-effects models are shown as text, with individual group comparisons indicated by asterisks: *: *P* < 0.05, **: P < 0.01, ***: *P* < 0.001.

Glucose provision did not significantly affect total leukocyte counts during infection, but a higher proportion of lymphocytes relative to neutrophils was observed in the 2.5% glucose group (Figure 2E–G). Glucose provision did, however, affect the plasma inflammatory response as levels of interleukin 10 (IL-10) were higher in the RES-SE group while levels of IL-6 were lower (Figure 2H,I). Alone, the SE infection had marked effects on the haematological and inflammatory response (Figure 2E-I). Finally, reduced glucose provision led to significantly lower plasma levels of AST and GGT, while BUN levels were elevated (Table 1). SE infection itself was associated with decreased levels of ALT, creatine kinase, and cholesterol, along with increased levels of inorganic phosphorus (Table 1).

**Table 1:** Plasma biochemistry profile at euthanasia.

|  | CON |  | P | SE |  | P | P<sub>SE</sub> |
| --- | --- | --- | --- | --- | --- | --- | --- |
|  | CON<br>(n = 3) | RES<br>(n = 4) |  | CON<br>(n = 13) | RES<br>(n = 13) |  |  |
| Alkaline Phosphatase | 295 [223 - 411] | 302 [192 - 413] | NS | 287 [204 - 400] | 316 [264 - 342] | NS | NS |
| ALT | 16.0 [12.0 - 19.0] | 13.8 [12.0 - 15.5] | NS | 12.2 [10.0 - 14.0] | 10.5 [10.0 - 11.0] | NS | <0.01 |
| AST | 40.3 [28.0 - 54.0] | 29.3 [18.5 - 40.0] | NS | 43.3 [12.0 - 55.0] | 31.2 [15.0 - 37.0] | 0.06 | NS |
| Albumin | 8.96 [8.04 - 9.45] | 9.16 [8.35 - 9.97] | NS | 8.52 [7.75 - 9.51] | 8.58 [7.28 - 9.58] | NS | NS |
| Creatin Kinase | 438 [390 - 479] | 622 [370 - 875] | NS | 335 [238 - 427] | 305 [264 - 329] | NS | <0.01 |
| Cholesterol | 1.72 [1.43 - 1.94] | 2.24 [1.93 - 2.54] | 0.08 | 1.45 [1.31 - 1.58] | 1.50 [1.42 - 1.62] | NS | <0.001 |
| Creatinine | 59.7 [56.0 - 62.0] | 59.8 [57.5 - 62.0] | NS | 63.1 [56.0 - 71.0] | 64.2 [56.0 - 71.0] | NS | NS |
| GGT | 31.0 [27.0 - 36.0] | 19.8 [14.5 - 25.0] | NS | 31.3 [25.0 - 39.0] | 24.5 [16.0 - 27.0] | <0.001 | NS |
| Inorganic Phosphorus | 1.40 [1.30 - 1.50] | 1.58 [1.40 - 1.75] | 0.08 | 1.83 [1.50 - 1.90] | 2.02 [1.90 - 2.3] | NS | <0.05 |
| Sodium | 142 [140 - 144] | 140 [138 - 142] | NS | 142 [139 - 145] | 138 [136 - 141] | NS | NS |
| Billirubin | 3.00 [3.00 - 3.00] | 3.25 [3.00 - 3.50] | NS | 3.38 [3.00 - 3.00] | 3.08 [3.00 - 3.00] | NS | NS |
| Total Protein | 20.8 [17.3 - 23.5] | 20.2 [18.3 - 22.1] | NS | 17.8 [15.8 - 21.6] | 18.3 [18.0 - 21.5] | NS | NS |
| Triglycerides | 2.00 [0.79 - 3.77] | 3.53 [2.29 - 4.77] | NS | 2.70 [0.79 - 3.93] | 2.89 [1.09 - 2.65] | NS | NS |
| BUN | 3.13 [2.50 - 4.00] | 5.85 [5.55 - 6.15] | <0.05 | 3.01 [2.80 - 3.20] | 6.01 [5.10 - 6.90] | <0.01 | NS |

Blood biochemical profile in newborn pigs infected with *Staphylococcus epidermidis* (SE) or saline control (CON), nourished with standard (CON) or restricted glucose (RES) PN. Shown as means with corresponding interquartile range. ALT: Alanine aminotransferase, AST: Aspartate aminotransferase, GGT: Gamma-glutamyl transferase. BUN: Blood urea nitrogen, NS: Not significant

We next assessed how glucose provision influenced host defence during infection. Blood bacterial densities did not differ between RES-SE and CON-SE groups at any time point (Sup. Fig. S1A). Using blood pH as a proxy for health, reaction norm analyses showed no correlation with bacterial load at 3 h in either group. At 6 h, a significant negative slope appeared in the RES-SE group only, suggesting activation of resistance mechanisms. By 12 h, this pattern reversed, with a negative slope in the CON-SE group. At euthanasia, no correlation was seen in either group (Sup. Fig. S1B).

Overall, the clinical and systemic immune and biochemical responses as well as defence strategies to both glucose reduction and SE infection were consistent with findings previously observed in this experimental model.

### Cardiac response to infection minimally affected by glucose provision

Next, we examined how glucose provision and SE infection influenced cardiac responses in the piglets. Relative heart weight was not affected by either glucose supply or SE infection (Figure 3A). Likewise, plasma troponin T levels did not differ between the CON or RES groups, nor between SE-infected and control animals (Figure 3B). Histological examination of formalin-fixed cardiac tissue stained with haematoxylin and eosin revealed no signs of necrosis or morphological abnormalities in any of the hearts. Neutrophilic infiltrates were observed in only three SE-infected animals and occurred at similar frequencies in the CON-SE (2/13) and RES-SE (1/13) groups (Sup. Figure 2A). Cardiomyocyte mitotic density and index was reduced in SE infected animals, but among the infected animals the mitotic density and index was higher in the RES-SE group (Figure 3C,D). There was no impact of SE infection or glucose provision on mitotic density in noncardiomyocytes (Sup. Figure 2B,C)

**Figure 3:**
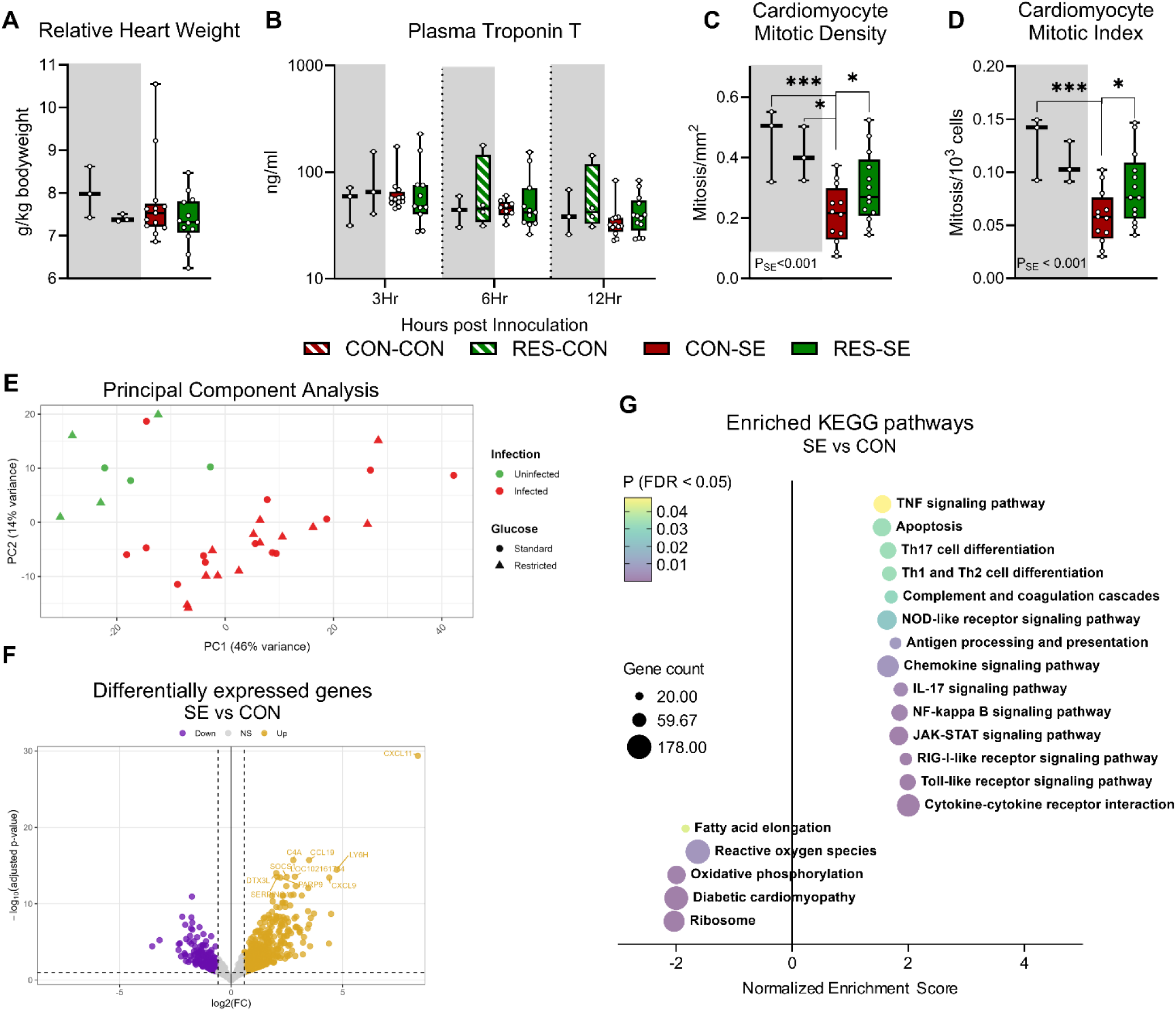
Cardiac responses in preterm pigs nourished with either standard (CON) or restricted (RES) glucose parenteral nutrition (PN) and infused with either *Staphylococcus epidermidis* (SE) bacteria or saline (CON). **A:** Heart weight relative to body weight. **B:** Plasma troponin levels at 3, 6, and 12 hours post-infection. **C:** Cardiomyocyte mitotic density across left and right ventricle, expressed as number of mitotic nuclei per mm^2^ **D:** Cardiomyocyte mitotic index across left and right ventricle, expressed as number of mitotic nuclei per 1000 total nuclei. **E:** Principal component analysis plot of cardiac transcriptomic profiles. **F:** Volcano plot showing differentially expressed genes (DEGs) in SE-infected versus control animals. **G:** Kyoto Encyclopedia of Genes and Genomes (KEGG) pathway enrichment analysis comparing SE-infected and control pigs; Positive normalized enrichment scores denotes upregulated pathways in SE infected piglets and negative downregulated.

We then investigated how glucose provision and SE infection affected cardiac gene expression. In a principal component analysis of the cardiac gene transcripts, infected and uninfected animals formed separate clusters, while no obvious differences were observed between pigs on standard or restricted glucose PN (Figure 3E). Interestingly, one infected animal clustered together with the uninfected, this animal was also the first to reach humane endpoints for euthanasia (7 hours post infection). In the heart tissue, differential gene expression analysis between infected piglets receiving standard (CON-SE) and reduced (RES-SE) glucose PN identified only three significantly regulated genes at an FDR<0.05 threshold. Among these, *FRMD5* and *PWWP3A* were downregulated, while *CHGA* was upregulated in the reduced-glucose group (Sup. Figure S3A). No KEGG pathways were differentially regulated, but Gene ontology enrichment analysis of biological properties associated with these transcripts indicated upregulation of immune related pathways in RES-SE animals such as *leukocyte degranulation*, *defence against bacterium* and *myeloid cell activation* (Sup. Figure S3B). No broader transcriptional changes or enrichment of metabolic pathways were observed. However, differential expression analysis comparing all infected and non-infected piglets identified a total of 1,332 significantly regulated genes in heart tissue at a FDR<0.05 threshold. Among the most strongly upregulated transcripts were *CXCL11*, *CCL19*, *C4A*, *LY6H*, and *DTX3L,* all immune related genes. Top downregulated genes were *SELE, APLNR, TMEFF2, ESM1* and *PKIB,* related to endothelial function or metabolism (Figure 3F). Gene set enrichment analysis based on KEGG pathways showed significant enrichment of multiple immune-related signalling cascades, including *cytokine–cytokine receptor interaction*, *JAK– STAT, NF-κB* and *toll-like receptor signalling pathways* (Figure 3G). These results demonstrate a marked transcriptional response to infection in cardiac tissue, upregulating a broad range of immune and inflammatory pathways. In addition, several metabolic pathways were also significantly downregulated in infected versus non-infected hearts, most notably *oxidative phosphorylation* and *fatty acid elongation* (Figure 3F). Interestingly, the genes *FABP3, FABP4, FABP5*, and *SCD5* were all strongly downregulated (FDR-adjusted P<0.001, Sup. Fig. S4). These genes act downstream of peroxisome proliferator-activated receptor alpha (PPAR-α) signalling, and although this overall pathway did not reach statistical significance in the KEGG analysis (FDR-adjusted P=0.10), the coordinated downregulation of these genes suggests a reduction in PPAR-α activity and mitochondrial β-oxidation.

## Discussion

Hyperglycaemia in preterm neonates is well known to associate with poorer infection outcomes, such as mortality and hospitalization[12, 15, 17, 41]. However, whether hyperglycaemia exacerbates infection-related neonatal cardiac dysfunction has not been reported. In our experiments, we show that experimental neonatal sepsis leads to cardiac dysfunction, with a drop in MAP, upregulation of inflammatory pathways and downregulation of metabolic ones in cardiac tissue. Likewise, in this experiment and previously, we show that high glucose intake and hyperglycaemia exacerbate neonatal sepsis[6, 29]. Our findings also suggest that animals in the reduced glucose group were able to initiate *disease resistance* responses earlier than animals on standard glucose PN and later transition to a *disease tolerance* strategy. This pattern aligns with our previous observations that survival in this sepsis model depends on early *disease resistance* to reduce pathogen burden, followed by *disease tolerance* to limit tissue damage and preserve host integrity[49]. However, despite these differences in clinical outcomes and a shift towards systemic *disease tolerance*, cardiac tissue demonstrated remarkable resilience to reduction in glucose intake. Minimal histological and transcriptomic changes were observed, suggesting that the neonatal heart maintains its functional and structural integrity despite drastic differences in glucose availability during infection, at least during the short duration of our experiments.

In *Experiment 1*, our finding that neonatal sepsis leads to loss of MAP aligns well with the clinical findings during neonatal sepsis[39]. Elevated levels of cardiac troponin are common in septic preterm infants and correlate with worse echocardiographic measures of cardiac function[2, 35]. A systematic review has likewise found that plasma troponin levels correlate with increased mortality in septic neonates[52]. However, in *Experiment 2* we could not identify differences in the increases in troponin as a cardiac biomarker. However, we could only investigate samples collected at 3, 6 and 12 hours post infection, since the cardiac puncture at euthanasia likely would increase troponin levels artificially[31]. It is likely that this short duration was not enough for meaningful increases in troponin to occur. Although infected hearts showed no necrosis, no troponin rise and only sparse neutrophil infiltration, cardiomyocyte mitotic activity fell markedly within 20 h of infection. This suggests that cell-cycle activity is an early and sensitive cellular readout of the septic neonatal heart, preceding overt structural injury. The pig heart is well suited to detect this, unlike rodents, porcine cardiomyocytes retain mitotic activity for up to two months after birth[45], similarly to how cardiomyocyte proliferation also continues in human infants[28]. Preterm birth itself already lowers cardiomyocyte proliferation in large-animal models[19, 46], so an additional, infection-driven suppression during this window could reduce final cardiomyocyte counts and have consequences beyond the acute illness.

Despite limited structural pathology, infected hearts showed clear transcriptional activation of inflammatory pathways together with suppression of energy metabolism, particularly oxidative phosphorylation. This occurred without increased troponin or substantial immune-cell infiltration, indicating an early cardiac response preceding overt injury. In adult sepsis models, PPAR-α signalling is similarly suppressed, and cardiomyocyte-specific loss of PPAR-α worsens cardiac dysfunction[40, 54]. Consistent with this, several PPAR-α target genes were downregulated in our infected hearts[18], suggesting that impaired mitochondrial fatty acid metabolism may contribute alongside inflammation to cardiac dysfunction during neonatal sepsis. Because cardiomyocytes normally rely predominantly on mitochondrial oxidative phosphorylation, with fatty acid β-oxidation supplying most ATP[8, 32], we expected glucose restriction to alter cardiac metabolic gene expression. However, reduced glucose provision had little effect on these pathways. It was nevertheless associated with higher cardiomyocyte mitotic activity in infected animals. Although glucose uptake can support cardiomyocyte proliferation in uninfected neonatal hearts[10], the benefit of glucose restriction during infection may instead be indirect, through reduced systemic inflammation and oxidative stress rather than altered cardiac substrate availability.

We have previously shown that reduction in glucose intake during neonatal infections increases mitochondrial metabolism in the liver[6, 51]. We have likewise shown that hyperglycaemia exacerbates inflammation and metabolic dysregulation in both the brain and kidneys[53]. We had expected similar effects in cardiac tissue, leading to reduction in cardiac inflammation. However, since the heart is less metabolically flexible and more reliant on fatty acid β-oxidation, it is possible that reduced glucose provision had minor impact on cardiac metabolism. However, there might also be differences between cardiac responses to sepsis in neonates versus adults. In a mouse model of polymicrobial sepsis, neonatal and aged mice exhibited different forms of sepsis-induced cardiac remodelling[37]. Although our data did show overt transcriptional changes, the preserved cardiomyocyte proliferation without associated injury markers may reflect better cardiac adaptation to inflammation. In neonatal mice, sepsis-induced inflammatory mediators triggered cardiomyocyte mitochondrial stress and reduced cardiac output[7]. Supporting mitochondrial metabolism during sepsis improves cardiac function in several animal models[16, 22, 23], and mitochondrial transplantation has improved both cardiac outcomes and survival[27]. Together, these findings suggest that mitochondrial dysfunction is not only involved in septic cardiac injury but may also represent a therapeutic target, potentially through PPAR-α activation and enhanced fatty acid β-oxidation.

### Limitations

Preterm pigs closely model human preterm infants in immune development and infection responses[5, 30, 36] and allow controlled PN administration, making them well suited for translational studies of neonatal sepsis and nutrition[1]. Continuous hemodynamic monitoring and cardiac transcriptomics enabled detailed characterization of the acute response to infection. However, *Experiment 1* included only three animals, limiting generalizability and robust statistical inference despite repeated-measures analysis; these findings should therefore be considered exploratory. The shorter survival in *Experiment 1* also suggests that anaesthesia and ventilation may have influenced disease progression. In *Experiment 2*, the short observation period and moderate sample size may have limited detection of delayed or more subtle cardiac effects. Finally, transcriptomics alone cannot fully capture changes in mitochondrial function or energy metabolism. Longer-term studies integrating metabolomic and proteomic analyses are therefore needed to define the mechanisms underlying cardiac resilience during neonatal sepsis.

## Conclusion

In this translational preterm pig model, neonatal bacterial infection induced circulatory collapse accompanied by cardiac activation of inflammatory pathways and suppression of mitochondrial oxidative metabolism. Reduced glucose provision attenuated systemic inflammation and was associated with higher survival, but had little effect on cardiac histopathology, injury markers, or gene expression. Despite limited structural injury, infection reduced cardiomyocyte mitotic activity, which appeared to be preserved by glucose restriction. These findings suggest that the neonatal heart shows substantial structural resilience during acute sepsis despite marked transcriptional responses to infection.

## Acknowledgements

We thank the Comparative Pediatrics and Nutrition (COPE) group at the University of Copenhagen for their support with the animal experiments, sample collection, and laboratory analyses.

## Consent

No human data reported in this paper. The Danish Animal Experiments Inspectorate approved all experimental procedures (license number 2020-15-0201-00520), in accordance with EU Directive 2010/63, governing the use of animals in research.

## Author contributions

OB, EM and DNN conceived the research questions, OB and TT performed the animal experiment, EM, JG and LO performed laboratory analyses. EM and OB performed statistical analyses. PL performed histological evaluation. OB drafted the paper, and all authors approved the final version.

## References

1. Aasmul-Olsen K, Pranevicius A, Klabunde B, Bæk O, Nguyen DN (2026) A Translational Preterm Piglet Model For Neonatal Sepsis. J Vis Exp. doi: 10.3791/71456

2. Abdel-Hady HE, Matter MK, El-Arman MM (2012) Myocardial dysfunction in neonatal sepsis: a tissue Doppler imaging study. Pediatr Crit Care Med 13:318–323. doi: 10.1097/PCC.0B013E3182257B6B

3. Alaedeen DI, Walsh MC, Chwals WJ (2006) Total parenteral nutrition–associated hyperglycemia correlates with prolonged mechanical ventilation and hospital stay in septic infants. J Pediatr Surg 41:239–244. doi: 10.1016/J.JPEDSURG.2005.10.045

4. Ayres JS (2020) The Biology of Physiological Health. Cell 181:250–269. doi: 10.1016/J.CELL.2020.03.036

5. Bæk O, Brunse A, Nguyen DN, Moodley A, Thymann T, Sangild PT (2020) Diet Modulates the High Sensitivity to Systemic Infection in Newborn Preterm Pigs. Front Immunol 11:1019. doi: 10.3389/fimmu.2020.01019

6. Bæk O, Muk T, Wu Z, Ye Y, Khakimov B, Casano AM, Gangadharan B, Bilic I, Brunse A, Sangild PT, Nguyen DN (2024) Altered hepatic metabolism mediates sepsis preventive effects of reduced glucose supply in infected preterm newborns. Elife 13. doi: 10.7554/ELIFE.97830.2

7. Denning NL, Aziz M, Diao L, Prince JM, Wang P (2020) Targeting the eCIRP/TREM-1 interaction with a small molecule inhibitor improves cardiac dysfunction in neonatal sepsis. Molecular Medicine 26:1–14. doi: 10.1186/S10020-020-00243-6/FIGURES/8

8. Depré C, Rider MH, Hue L (1998) Mechanisms of control of heart glycolysis. Eur J Biochem 258:277–290. doi: 10.1046/J.1432-1327.1998.2580277.X

9. van Diest PJ, Baak JPA, Matze-Cok P, Wisse-Brekelmans ECM, van Galen CM, Kurver PHJ, Bellot SM, Fijnheer J, van Gorp LHM, Kwee WS, Los J, Peterse JL, Ruitenberg HM, Schapers RFM, Schipper MEI, Somsen JG, Willig AWPM, Ariens AT (1992) Reproducibility of mitosis counting in 2,469 breast cancer specimens: Results from the Multicenter Morphometric Mammary Carcinoma Project. Hum Pathol 23:603–607. doi: 10.1016/0046-8177(92)90313-R

10. Fajardo VM, Feng I, Chen BY, Perez-Ramirez CA, Shi B, Clark P, Tian R, Lien CL, Pellegrini M, Christofk H, Nakano H, Nakano A (2021) GLUT1 overexpression enhances glucose metabolism and promotes neonatal heart regeneration. Sci Rep 11. doi: 10.1038/S41598-021-88159-X

11. Graham PL, Begg MD, Larson E, Della-Latta P, Allen A, Saiman L (2006) Risk factors for late onset gram-negative sepsis in low birth weight infants hospitalized in the neonatal intensive care unit. Pediatric Infectious Disease Journal 25:113–117. doi: 10.1097/01.INF.0000199310.52875.10

12. Hall NJ, Peters M, Eaton S, Pierro A (2004) Hyperglycemia is associated with increased morbidity and mortality rates in neonates with necrotizing enterocolitis. J Pediatr Surg 39:898–901. doi: 10.1016/j.jpedsurg.2004.02.005

13. Harbeson D, Ben-Othman R, Amenyogbe N, Kollmann TR (2018) Outgrowing the Immaturity Myth: The Cost of Defending From Neonatal Infectious Disease. Front Immunol 9:1077. doi: 10.3389/fimmu.2018.01077

14. Harbeson D, Francis F, Bao W, Amenyogbe NA, Kollmann TR (2018) Energy Demands of Early Life Drive a Disease Tolerant Phenotype and Dictate Outcome in Neonatal Bacterial Sepsis. Front Immunol 9:1918. doi: 10.3389/fimmu.2018.01918

15. Hay WW, Rozance PJ (2018) Neonatal Hyperglycemia—Causes, Treatments, and Cautions. Journal of Pediatrics 200:6–8. doi: 10.1016/j.jpeds.2018.04.046

16. Hou D, Liao H, Hao S, Liu R, Huang H, Duan C (2024) Curcumin simultaneously improves mitochondrial dynamics and myocardial cell bioenergy after sepsis via the SIRT1-DRP1/PGC-1α pathway. Heliyon 10. doi: 10.1016/j.heliyon.2024.e28501

17. Kao LS, Morris BH, Lally KP, Stewart CD, Huseby V, Kennedy KA (2006) Hyperglycemia and morbidity and mortality in extremely low birth weight infants. J Perinatol 26:730–736. doi: 10.1038/SJ.JP.7211593

18. Kersten S, Rakhshandehroo M, Knoch B, Müller M (2010) Peroxisome Proliferator-Activated Receptor Alpha Target Genes. PPAR Res 2010:612089. doi: 10.1155/2010/612089

19. Kim MY, Eiby YA, Lumbers ER, Wright LL, Gibson KJ, Barnett AC, Lingwood BE (2014) Effects of glucocorticoid exposure on growth and structural maturation of the heart of the preterm piglet. PLoS One 9. doi: 10.1371/JOURNAL.PONE.0093407

20. Klabunde B, Bæk O, Aasmul-Olsen K, Hammerich Thysen A, Gudbrandsdottir M, Roed Spiegelhauer M, Laakmann K, Hoffmann K, Schmeck B, Brunse A, Long NP, Le Q-V, Brix S, Chawes B, Nguyen DN (2026) Ketone metabolism promotes disease tolerance and survival in experimental neonatal sepsis. Cell Communication and Signaling 2026. doi: 10.1186/S12964-026-03136-Z

21. Kung YH, Hsieh YF, Weng YH, Lien RI, Luo J, Wang Y, Huang YC, Chen CL, Chen CJ (2016) Risk factors of late-onset neonatal sepsis in Taiwan: A matched case-control study. Journal of Microbiology, Immunology and Infection 49:430–435. doi: 10.1016/J.JMII.2013.10.001

22. Lazarević M, Kostić M, Džopalić T, Sokolović D, Lazarević Z, Milovanović J, Ničković V, Sokolović D (2024) Melatonin Mediates Cardiac Tissue Damage under Septic Conditions Induced by Lipopolysaccharide. Int J Mol Sci 25:11088. doi: 10.3390/IJMS252011088

23. Liu H, Hu Q, Ren K, Wu P, Wang Y, Lv C (2023) ALDH2 mitigates LPS-induced cardiac dysfunction, inflammation, and apoptosis through the cGAS/STING pathway. Mol Med 29. doi: 10.1186/S10020-023-00769-5

24. Loftus RM, Finlay DK (2016) Immunometabolism: Cellular Metabolism Turns Immune Regulator. J Biol Chem 291:1–10. doi: 10.1074/jbc.R115.693903

25. Medzhitov R, Schneider DS, Soares MP (2012) Disease tolerance as a defense strategy. Science 335:936–941. doi: 10.1126/SCIENCE.1214935

26. Mesotten D, Joosten K, van Kempen A, Verbruggen S, Braegger C, Bronsky J, Cai W, Campoy C, Carnielli V, Darmaun D, Decsi T, Domellöf M, Embleton N, Fewtrell M, Fidler Mis N, Franz A, Goulet O, Hartman C, Hill S, Hojsak I, Iacobelli S, Jochum F, Joosten K, Kolaček S, Koletzko B, Ksiazyk J, Lapillonne A, Lohner S, Mesotten D, Mihályi K, Mihatsch WA, Mimouni F, Mølgaard C, Moltu SJ, Nomayo A, Picaud JC, Prell C, Puntis J, Riskin A, Saenz De Pipaon M, Senterre T, Shamir R, Simchowitz V, Szitanyi P, Tabbers MM, Van Den Akker CHB, Van Goudoever JB, Van Kempen A, Verbruggen S, Wu J, Yan W (2018) ESPGHAN/ESPEN/ESPR/CSPEN guidelines on pediatric parenteral nutrition: Carbohydrates. Clin Nutr 37:2337–2343. doi: 10.1016/J.CLNU.2018.06.947

27. Mokhtari B, Hamidi M, Badalzadeh R, Mahmoodpoor A (2023) Mitochondrial transplantation protects against sepsis-induced myocardial dysfunction by modulating mitochondrial biogenesis and fission/fusion and inflammatory response. Mol Biol Rep 50:2147–2158. doi: 10.1007/S11033-022-08115-4

28. Mollova M, Bersell K, Walsh S, Savla J, Das LT, Park SY, Silberstein LE, Dos Remedios CG, Graham D, Colan S, Kühn B (2013) Cardiomyocyte proliferation contributes to heart growth in young humans. Proc Natl Acad Sci U S A 110:1446– 1451. doi: 10.1073/PNAS.1214608110/-/DCSUPPLEMENTAL/SM06.MOV

29. Muk T, Brunse A, Henriksen NL, Aasmul-Olsen K, Nguyen DN (2022) Glucose supply and glycolysis inhibition shape the clinical fate of Staphylococcus epidermidis– infected preterm newborns. JCI Insight 7. doi: 10.1172/JCI.INSIGHT.157234

30. Nguyen DN, Jiang P, Frøkiær H, Heegaard PMH, Thymann T, Sangild PT (2016) Delayed development of systemic immunity in preterm pigs as a model for preterm infants. Sci Rep 6:36816. doi: 10.1038/srep36816

31. O’Brien PJ, Smith DEC, Knechtel TJ, Marchak MA, Pruimboom-Brees I, Brees DJ, Spratt DP, Archer FJ, Butler P, Potter AN, Provost JP, Richard J, Snyder PA, Reagan WJ (2006) Cardiac troponin I is a sensitive, specific biomarker of cardiac injury in laboratory animals. Lab Anim 40:153–171. doi: 10.1258/002367706776319042

32. Onay-Besikci A (2006) Regulation of cardiac energy metabolism in newborn. Mol Cell Biochem 287:1–11. doi: 10.1007/S11010-006-9123-9

33. O’Neill LAJ, Kishton RJ, Rathmell J (2016) A guide to immunometabolism for immunologists. Nat Rev Immunol 16:553–565. doi: 10.1038/nri.2016.70

34. Purisch SE, Gyamfi-Bannerman C (2017) Epidemiology of preterm birth. Semin Perinatol 41:387–391. doi: 10.1053/J.SEMPERI.2017.07.009

35. Ramadhina N, Sukardi R, Advani N, Rohsiswatmo R, Putra ST, Djer MM (2015) Ventricular function and high-sensitivity cardiac troponin T in preterm infants with neonatal sepsis. Paediatr Indones 55:203–8. doi: 10.14238/PI55.4.2015.203-8

36. Ren S, Hui Y, Obelitz-Ryom K, Brandt AB, Kot W, Nielsen DS, Thymann T, Sangild PT, Nguyen DN (2018) Neonatal gut and immune maturation is determined more by postnatal age than by post-conceptional age in moderately preterm pigs. Am J Physiol Gastrointest Liver Physiol. doi: 10.1152/ajpgi.00169.2018

37. Sanchez KK, McCarville JL, Stengel SJ, Snyder JM, Williams AE, Ayres JS (2026) Disease tolerance and infection pathogenesis age-related tradeoffs in mice. Nature 650:727–735. doi: 10.1038/S41586-025-09923-X

38. Schneider DS, Ayres JS (2008) Two ways to survive infection: what resistance and tolerance can teach us about treating infectious diseases. Nat Rev Immunol 8:889–895. doi: 10.1038/NRI2432;KWRD

39. Shane AL, Sánchez PJ, Stoll BJ (2017) Neonatal sepsis. The Lancet 390:1770–1780. doi: 10.1016/S0140-6736(17)31002-4

40. Standage SW, Bennion BG, Knowles TO, Ledee DR, Portman MA, McGuire JK, Liles WC, Olson AK (2017) PPARα augments heart function and cardiac fatty acid oxidation in early experimental polymicrobial sepsis. Am J Physiol Heart Circ Physiol 312:H239–H249. doi: 10.1152/AJPHEART.00457.2016

41. Stensvold HJ, Strommen K, Lang AM, Abrahamsen TG, Steen EK, Pripp AH, Ronnestad AE (2015) Early Enhanced Parenteral Nutrition, Hyperglycemia, and Death Among Extremely Low-Birth-Weight Infants. JAMA Pediatr 169:1003–1010. doi: 10.1001/JAMAPEDIATRICS.2015.1667

42. Strunk T, Molloy EJ, Mishra A, Bhutta ZA (2024) Neonatal bacterial sepsis. The Lancet 404:277–293. doi: 10.1016/S0140-6736(24)00495-1

43. Tröger B, Göpel W, Faust K, Müller T, Jorch G, Felderhoff-Müser U, Gortner L, Heitmann F, Hoehn T, Kribs A, Laux R, Roll C, Emeis M, Mögel M, Siegel J, Vochem M, von der Wense A, Wieg C, Herting E, Härtel C, German Neonatal Network (2014) Risk for late-onset blood-culture proven sepsis in very-low-birth weight infants born small for gestational age: a large multicenter study from the German Neonatal Network. Pediatr Infect Dis J 33:238–43. doi: 10.1097/INF.0000000000000031

44. Uthaya S, Modi N (2014) Practical preterm parenteral nutrition: Systematic literature review and recommendations for practice. Early Hum Dev 90:747–753. doi: 10.1016/j.earlhumdev.2014.09.002

45. Velayutham N, Alfieri CM, Agnew EJ, Riggs KW, Baker RS, Ponny SR, Zafar F, Yutzey KE (2020) Cardiomyocyte cell cycling, maturation, and growth by multinucleation in postnatal swine. J Mol Cell Cardiol 146:95–108. doi: 10.1016/j.yjmcc.2020.07.004

46. Vrselja A, Pillow JJ, Bensley JG, Ellery SJ, Ahmadi-Noorbakhsh S, Moss TJ, Black MJ (2022) Intrauterine inflammation exacerbates maladaptive remodeling of the immature myocardium after preterm birth in lambs. Pediatr Res 92:1555–1565. doi: 10.1038/S41390-022-01955-7

47. Webbe J, Battersby C, Longford N, Oughham K, Uthaya S, Modi N, Gale C (2022) Use of parenteral nutrition in the first postnatal week in England and Wales: an observational study using real-world data. BMJ Paediatr Open 6. doi: 10.1136/BMJPO-2022-001543,

48. Weis S, Carlos AR, Moita MR, Singh S, Blankenhaus B, Cardoso S, Larsen R, Rebelo S, Schäuble S, Del Barrio L, Mithieux G, Rajas F, Lindig S, Bauer M, Soares MP (2017) Metabolic Adaptation Establishes Disease Tolerance to Sepsis. Cell 169:1263–1275.e14. doi: 10.1016/J.CELL.2017.05.031

49. Wu Z, Bæk O, Muk T, Yang L, Shen RL, Gangadharan B, Bilic I, Nielsen DS, Sangild PT, Nguyen DN (2024) Feeding cessation and antibiotics improve clinical symptoms and alleviate gut and systemic inflammation in preterm pigs sensitive to necrotizing enterocolitis. Biomedicine and Pharmacotherapy 179. doi: 10.1016/j.biopha.2024.117391

50. Wu Z, Tien NTN, Bæk O, Zhong J, Klabunde B, Nguyen TT, Yen NTH, Long NP, Nguyen DN (2024) Regulation of host metabolism and defense strategies to survive neonatal infection. Biochim Biophys Acta Mol Basis Dis 1870. doi: 10.1016/J.BBADIS.2024.167482

51. Wu Z, Tran N, Tien N, Klabunde B, Aasmul-Olsen K, Offersen SM, Thi N, Yen H, Muk T, Thysen AH, Brix S, Brustad N, Wang T, Stokholm J, Bønnelykke K, Brunse A, Phuoc Long N, Chawes B, Baek O, Duc ✉, Nguyen N (2026) Harnessing the glycolysis-TCA cycle axis to boost host defense against neonatal infection. EMBO Molecular Medicine 2026 1–24. doi: 10.1038/S44321-026-00463-Z

52. Zheng P, Wang X, Guo T, Gao W, Huang Q, Yang J, Gao H, Liu Q (2023) Cardiac troponin as a prognosticator of mortality in patients with sepsis: A systematic review and meta-analysis. Immun Inflamm Dis 11:e1014. doi: 10.1002/IID3.1014

53. Zhong J, Bæk O, Doughty R, Jørgensen BM, Jensen HE, Thymann T, Sangild PT, Brunse A, Nguyen DN (2025) Reduced parenteral glucose supply during neonatal infection attenuates neurological and renal pathology associated with modulation of innate and Th1 immunity. Biochim Biophys Acta Mol Basis Dis 1871. doi: 10.1016/J.BBADIS.2025.167723

54. Zhu X Xin, Wang X, Jiao S Yu, Liu Y, Shi L, Xu Q, Wang J Jing, Chen Y Er, Zhang Q, Song Y Ting, Wei M, Yu B Ai, Fielitz J, Gonzalez FJ, Du J, Qu A Juan (2023) Cardiomyocyte peroxisome proliferator-activated receptor α prevents septic cardiomyopathy via improving mitochondrial function. Acta Pharmacol Sin 44:2184. doi: 10.1038/S41401-023-01107-5

