## Supplemental Figures for "Glucose restriction reduces systemic, but not cardiac, inflammation during neonatal sepsis in preterm pigs"

### Supplementary Materials

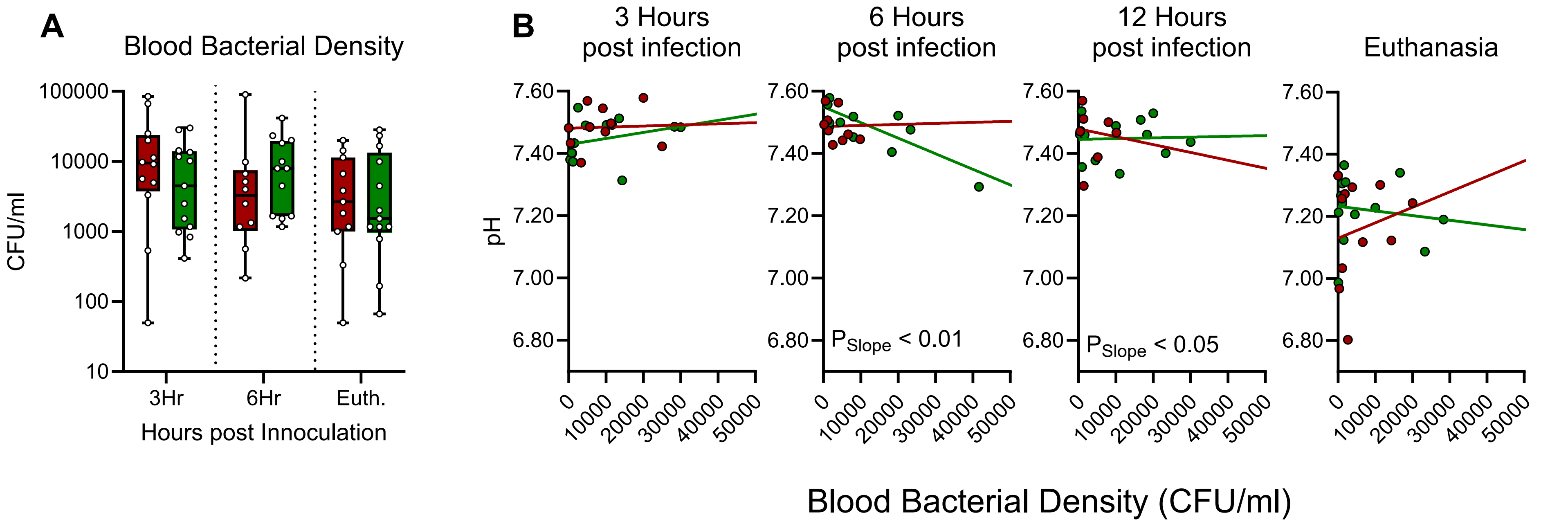


**Supplementary Figure S2:** Pathogen burden and evaluation of host defense strategies in preterm pigs infected with *Staphylococcus epidermidis* (SE) and nourished with either standard (CON) or restricted (RES) glucose parenteral nutrition (PN). **A:** Blood bacterial density over the course of the experiment. **B:** Reaction norm analyses at 3, 6, and 12 hours post-infection and at euthanasia, showing the relationship between blood bacterial density and blood pH. $P_{slope}$ indicates a significant difference in the slopes between the two treatment groups.

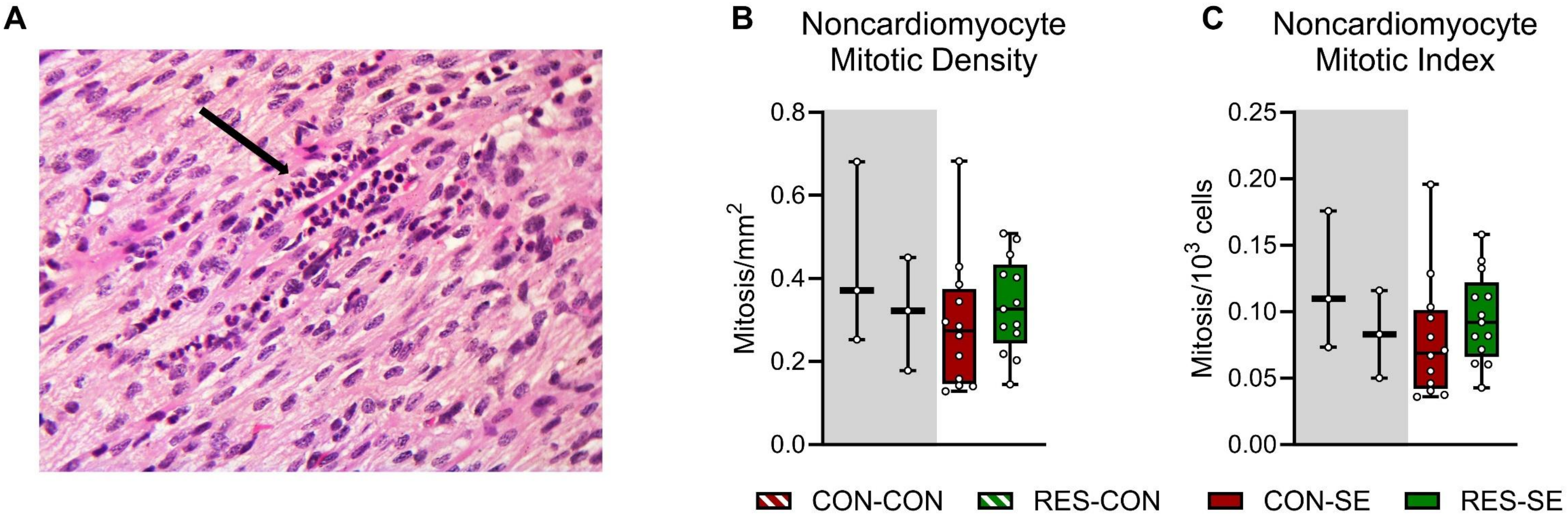


**Supplementary Figure S2: A:** Representative microscopy image of hematoxylin and eosin (H&E)-stained cardiac tissue from an SE-infected preterm pig, with arrow indicating neutrophil infiltration. **B:** Noncardiomyocyte mitotic density across left and right ventricle, expressed as number of mitotic nuclei per $mm^2$ **C:** Noncardiomyocyte mitotic index across left and right ventricle, expressed as number of mitotic nuclei per 1000 total nuclei.

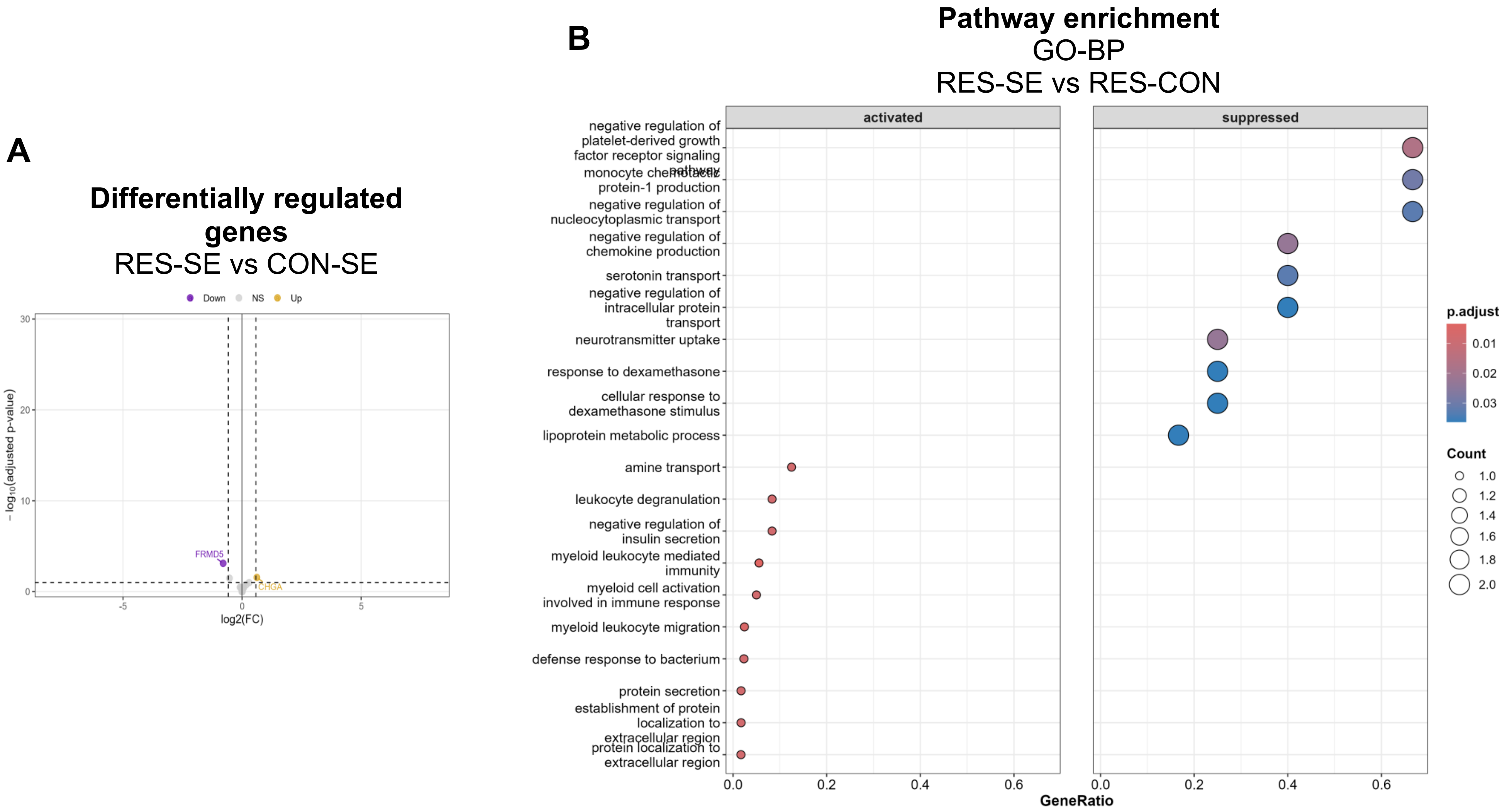


**Supplementary Figure S3.** Supplemental gene expression results. **A:** Volcano plot showing differentially expressed genes (DEGs) in *Staphylococcus epidermidis* (SE)-infected preterm pigs receiving restricted (RES) versus standard (CON) glucose parenteral nutrition. **B:** Pathway enrichment analysis based on Gene Ontology – Biological Process (GO-BP) terms in SE-infected RES versus CON animals.

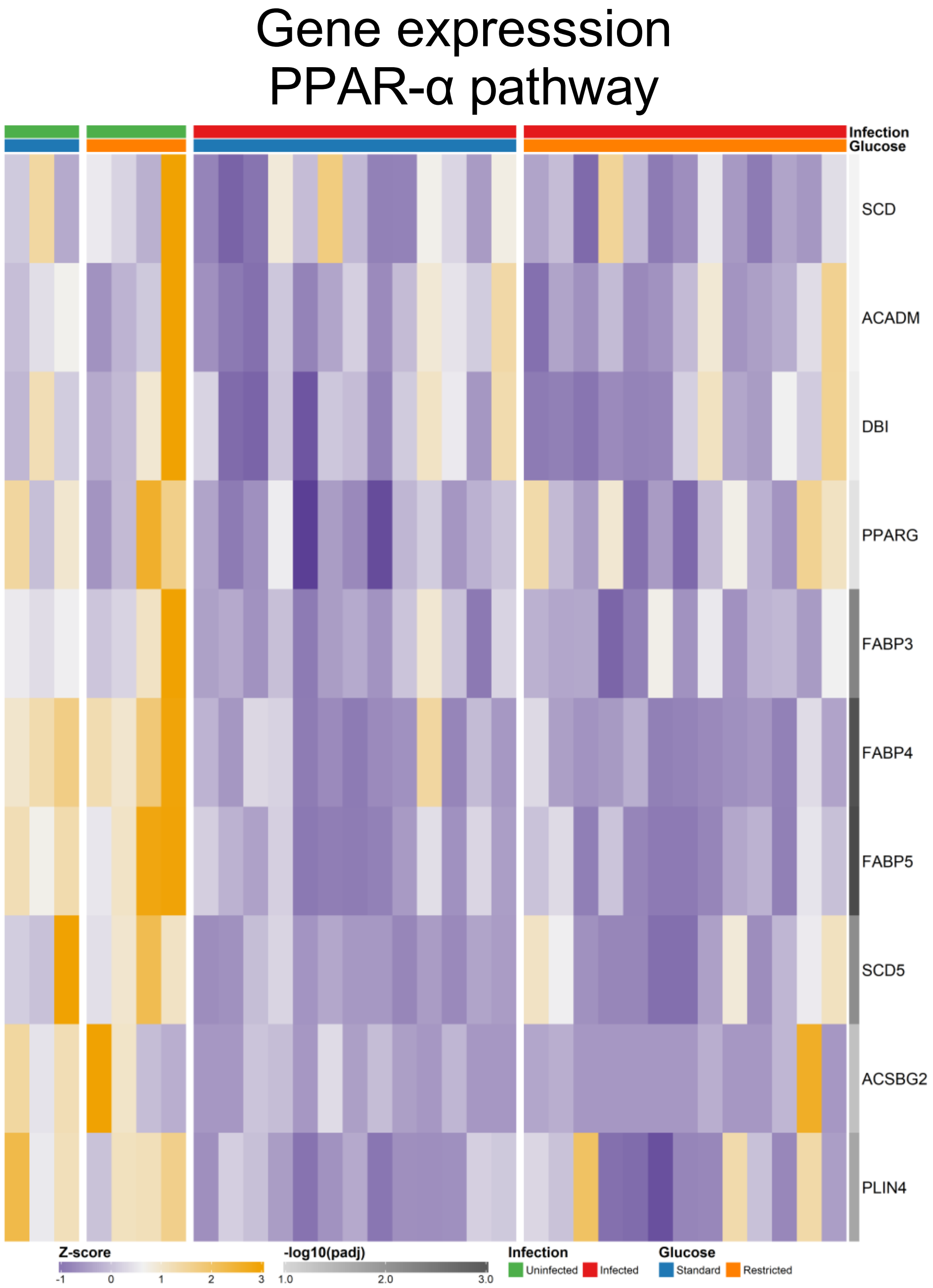


**Supplementary Figure 4.** Heatmap showing expression of genes within the peroxisome proliferator-activated receptor alpha (PPAR-α) pathway, with yellow indicating high expression and purple indicating low expression.
